# Optimization of cryo-electron microscopy for quantitative analysis of lipid bilayers

**DOI:** 10.1101/2022.08.23.505005

**Authors:** Frederick A. Heberle, Doug Welsch, Haden L. Scott, M. Neal Waxham

## Abstract

Cryogenic electron microscopy (cryo-EM) is among the most powerful tools available for interrogating nanoscale structure of biological structures. We recently showed that cryo-EM can be used to measure the bilayer thickness of lipid vesicles and biological membranes with sub-angstrom precision, resulting in the direct visualization of nanoscopic domains of different thickness in multicomponent lipid mixtures and giant plasma membrane vesicles. Despite the great potential of cryo-EM for revealing the lateral organization of biomembranes, a large parameter space of experimental conditions remains to be optimized. Here, we systematically investigate the influence of instrument parameters and image post-processing steps on the ability to accurately measure bilayer thickness and discriminate regions of different thickness within unilamellar liposomes. We also demonstrate a spatial autocorrelation analysis to extract additional information about lateral heterogeneity.

**Significance:** Raft domains in unstimulated cells have proven difficult to directly visualize owing to their nanoscopic size and fleeting existence. The few techniques capable of nanoscopic spatial resolution typically rely on interpretation of indirect spectroscopic or scattering signals or require stabilizing the membrane on a solid support. In contrast, cryo-EM yields direct images of nanoscale domains in probe-free, unsupported membranes. Here, we systematically optimize key steps in the experimental and analysis workflow for this new and specialized application. Our findings represent an important step toward developing cryo-EM into a robust method for investigating phase behavior of membranes at length scales relevant to lipid rafts.

## Introduction

Cell membranes have a remarkable capacity for self-organization conferred by the structural diversity of their lipidomes (1). It is increasingly clear that within the outermost plasma membrane (PM), nonideal interactions between different classes of lipids results in clustering or even phase separation that can in turn influence the spatial organization of proteins (2). The membrane raft hypothesis, which grew from the work of Kai Simons and co-workers (3), has triggered extensive research to uncover roles for membrane microdomains in cell functions including protein trafficking and cell signaling (4). Rafts in cells are transient and nanoscopic in size but can be induced to coalesce into larger platforms upon stimulation, for example the crosslinking of immune receptors on the cell surface (5). Simplified model systems composed of a few representative lipids have been invaluable for teasing apart the physicochemical underpinnings of these self-organizing phenomena (6). These efforts have culminated in a consensus that domain formation is driven primarily by unfavorable interactions between intrinsically ordered (e.g., sphingomyelin) and disordered (e.g., unsaturated phosphocholine) lipids, while the presence of cholesterol ensures that ordered domains are fluid (7).

Techniques sensitive to phase separation in membranes can generally be divided into those providing indirect evidence of nonrandom mixing and those that provide direct (i.e., visual) evidence. Among indirect techniques, ^2^H-NMR (8) or ESR spectra (9) can in favorable cases be decomposed into signal arising from two distinct environments, while abrupt changes in FRET between donor and acceptor lipids can occur when phase separation causes spatial reorganization of the probes (10)(11)(12). With specialized contrast-matching schemes, SANS is sensitive to phase separation of protiated and deuterated lipids (13)(14). All indirect methods ultimately involve interpreting an ensemble-averaged signal of limited information content, and most depend on some type of model fitting. In contrast, direct visualization by AFM (15), secondary ion mass spectrometry imaging (SIMS) (16)(17), or one of the various super-resolution microscopies (18) can in principle allow for detailed information about domain size distributions, though in practice this goal has not yet been achieved. A significant drawback of AFM and SIMS is the requirement of a solid support that can perturb lipid phase behavior (19)(20). While super-resolution microscopies are rapidly improving (21), few of the existing modalities are truly capable of < 20 nm resolution, and each of these (i.e., PALM, STORM, NSOM) suffers from slow image acquisition time that blurs information in the spatial regime (18).

The advent of cryogenic electron microscopy (cryo-EM) has revolutionized the field of structural biology, allowing for near-atomic resolution structures of proteins and nucleic acids (22)(23)(24). Specimens for cryo-EM are prepared by plunging an aqueous sample of a biological material of interest into liquid ethane (–188°C) to rapidly vitrify the sample and lock in the ensemble of structures found under native conditions. Vitreous ice lacks long-range structure, a fact that is crucial both for preserving the sample and for obtaining high quality EM images (hexagonal ice diffracts electrons strongly). Importantly, image contrast in cryo-EM does not rely on the addition of any exogenous probe or electron-dense stains that can perturb the native lipid organization.

While many cryo-EM studies have used lipid vesicles as a means for imaging reconstituted membrane proteins (25)(26), few have focused on the membranes themselves. Two such studies have paved the way for our own work. Tahara et al. showed that the spacing between the two dark concentric circles seen in projection images of lipid vesicles increases with increasing acyl chain length, thus demonstrating sensitivity to membrane thickness (27). Sigworth and co-workers used atomistic MD simulations to calculate cryo-EM projections of lipid vesicles which, when combined with experimental images, provided compelling evidence that the membrane dipole potential contributes significantly to its electron scattering profile (28). Building on these earlier works, we (simultaneously with the Keller group) were the first to demonstrate that the inherent thickness differences between ordered and disordered phases provides sufficient contrast to resolve coexisting domains within individual vesicles in cryo-EM images (29)(30). While these proof-of-principle studies establish the potential of cryo-EM for analyzing membrane structure and function, a vast parameter space remains to be optimized, and specialized analysis tools developed, before this potential can be fully realized.

To this end, we have undertaken a systematic optimization of key steps in the experimental and analysis workflow for extracting membrane thickness from projection images of single-component and multi-component lipid vesicles. We report the influence on this measurement of instrument parameters such as electron dose, defocus length, and energy filtering, as well as post-processing steps including high pass filtering and contrast transfer function corrections. We also demonstrate an autocorrelation analysis of the spatially-resolved bilayer thickness that can in principle provide estimates of domain sizes.

## Materials and methods

### Chemicals

1,2-dioleoyl-*sn*-glycero-3-phosphocholine (DOPC), 1,2-dipalmitoyl-*sn*-glycero-3-phosphocholine (DPPC), 1-palmitoyl-2-oleoyl-*sn*-glycero-3-phospho-(1’-rac-glycerol) sodium salt (POPG), and cholesterol (chol) were purchased from Avanti Polar Lipids (Alabaster, AL) as dry powders and used as supplied. Stock solutions of phospholipids and cholesterol were prepared by weighing powder directly into a volumetric flask and dissolving in HPLC-grade chloroform; two tightly-bound water molecules per phospholipid were assumed present in calculations of molar concentrations. Stocks were stored at −80°C until use. Ultrapure H_2_O was obtained from a Milli-Q IC 7000 purification system (Millipore Sigma, Burlington, MA).

### Preparation and cryopreservation of large unilamellar vesicles

All data in this study was collected on either single-phase vesicle preparations composed of DOPC/POPG (95/5%) or phase-separated vesicles composed of DPPC/DOPC/POPG/chol (40/35/5/20%). Aqueous lipid dispersions at 3 mg/mL total lipid concentration were prepared by first mixing appropriate volumes of lipid stocks in chloroform with a glass Hamilton syringe. The solvent was evaporated with an inert gas stream followed by vacuum desiccation overnight. The dry lipid film was hydrated with ultrapure water at 45°C for at least 1 h with intermittent vortex mixing. The resulting multilamellar vesicle (MLV) suspension was subjected to at least 5 freeze/thaw cycles between a −80°C freezer and a 45°C water bath, and was then extruded through a 0.1 μm polycarbonate filter using a handheld mini-extruder (Avanti Polar Lipids) by passing the suspension through the filter 31 times. DOPC/POPG vesicles were extruded at room temperature, and DPPC/DOPC/POPG/chol vesicles were extruded at 45°C (~10°C above the miscibility transition temperature for this mixture). The size and polydispersity of each vesicle preparation was assessed using dynamic light scattering immediately after preparation and immediately before cryo-preservation, which was typically performed 2-3 days after preparation.

To cryopreserve vesicles, 4 μL of sample were applied to a Quantifoil 2/2 carbon-coated 200 mesh copper grid that was glow-discharged for 30 sec at 20 mA in a Pelco Easi-Glow discharge device. After manual blotting at room temperature (~22°C), the grids were plunged into liquid ethane cooled with liquid N2. Cryo-preserved grids were stored in liquid N2 until use.

### Image acquisition

Image collection was accomplished at various values of underfocus on a Thermo Fisher Titan Krios operated at 300 keV equipped with a Gatan K2 Summit direct electron detector operated in counting mode. Where noted, data was collected with a Gatan BioQuantum energy filter with a 20 eV slit in zero loss mode. Data collection was performed in a semi-automated fashion using SerialEM software operated in low-dose mode (31). Briefly, atlases were produced in the EPU software, and then areas of interest with appropriate ice thickness were identified. Then, 8×8 montages were collected at low magnification (4800×) at various positions across the grid, with individual areas marked for automated data collection. Data was collected at 53,000× nominal magnification equating to 2.66 Å/pixel. Nine-second movies of 30 dose-fractionated frames (0.3 sec/frame) were collected at each target site with the total electron dose kept to < 20 e^−^/Å^2^. For experiments investigating the effect of electron dose on the sample, the movie acquisition was extended to 60 sec. Using scripts in MotionCor2 (32) the movies were motion corrected for subsequent analysis. For the analysis of dose damage, the same signal-to-noise in each analyzed image was maintained by sequentially choosing the same number of frames for correction at 10-12 sec intervals of the 60 sec movie.

### Image selection and processing

The defocus/astigmatism of each image was assessed with CTFFind4 (v4.1.13) and images less than 1 μm or more than 4 μm underfocus were discarded. Typical image processing steps include high pass filtering to remove low-frequency variation in intensity (due to e.g. gradients in ice thickness) and phase-flipping to partially reverse the effects of the contrast transfer function. High pass filtering of images was accomplished with the *mtffilter* command in IMOD. Phase-flipping was accomplished taking parameters (defocus and azimuth of astigmatism) from CTFFind4 to create scripts for correction using the *ctfphaseflip* routine in IMOD. To determine the influence of various image processing steps on bilayer thickness, drift corrected images that were either high pass filtered, high pass filtered and phase-flipped, or neither high pass filtered nor phase-flipped (i.e., raw images) were analyzed.

### Calculation of the contrast transfer function

The contrast transfer function, *c*(***s***), used to generate the results shown in Fig. 5 is given by:

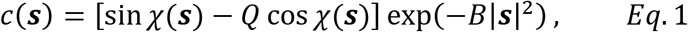

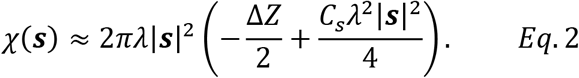

In Eqs. 1–2, ***s*** is the spatial frequency in units of Å^−1^, *Q* is the unitless amplitude contrast factor, *B* is the amplitude decay factor, *χ*(***s***) is the phase perturbation factor, *λ* is the electron wavelength, Δ*Z* is the defocus length, and *C*_*s*_ is the spherical aberration coefficient. For the calculations presented in Fig. 5 we used typical values from our experiments: *Q* =7.5%, *B* = 300 Å^2^, *λ* = 0.0197 Å (300 keV), and Δ*Z* ranging from 0-4 μm underfocus. We neglected the spherical aberration as the coefficient for our instrument (*C*_*s*_ = 2 mm) has an insignificant influence on the CTF.

### Data analysis and statistics

Individual vesicle contours (identifying the center of the lipid bilayer) from raw, high pass filtered, or high pass filtered/phase-flipped images were produced using a neural network-based learning algorithm (MEMNET) developed and kindly provided by Dr. Tristan Bepler (MIT). All subsequent analyses were performed using custom Mathematica v. 12.0 (Wolfram Research, Inc.) routines as previously described (30). Summary data from the Mathematica analyses were plotted and statistical analysis performed using Prism (v.9.4.1, Graphpad).

A key measurement obtained from the analysis is the spatially resolved bilayer thickness, details of which are provided in our previous work (30). Briefly, bilayer thickness is measured along the projected circumference of individual vesicles. The minimum spatial resolution *ρ* of the thickness measurement corresponds to an arc length over which the image intensity is radially averaged (for all data presented in this work, *ρ* was chosen to be 5 nm). The bilayer thickness *t* is then calculated as the distance between the two minima in the radially averaged intensity profile. For a vesicle of diameter *d*, the number of thickness measurements obtained in this way is *N* = ⌊*πd*/*ρ*⌋. We also computed the lag-*h* autocorrelation of spatially resolved bilayer thickness *ξ*_*h*_ for individual vesicles,

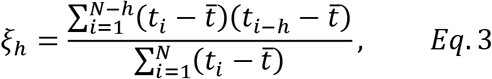

where *i* indexes the spatially ordered list of *N* thickness measurements *t*_*i*_ in vesicle *j* and 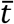 is the mean bilayer thickness averaged over all segments of all vesicles:

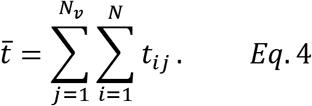

For individual vesicles, *ξ*_*h*_ was calculated for lags *h* = 1 to *h* = *N*/2. The set of lag-*h* values for all vesicles were then used to calculate an average *ξ*_*h*_ and associated uncertainty and converted to autocorrelation data *ξ*(*r*), where the lag distance *r* = *ρh*.

## Results and discussion

We and others have previously established the utility of cryo-EM for quantifying physical characteristics of synthetic bilayers (29)(26)(30)(33). Here, we investigated how instrument parameters and image processing can be tuned to optimize the information content of these experiments using two model systems: a mixture of DOPC/POPG (95/5) that is uniformly mixed at room temperature, and a ternary mixture of DPPC/DOPC/POPG/chol (40/35/5/20%) that exhibits coexisting liquid phases (Ld+Lo) at room temperature. In both cases, we prepared extruded 100 nm vesicles using a small amount of negatively charged lipid (5 mol% POPG) to minimize vesicle aggregation and the occurrence of multi-bilayer vesicles (33).

Our results are organized as follows. First, we discuss the influence of instrument parameters (data acquisition stage) on measurements. Second, we discuss the influence of post-acquisition image processing. Finally, we demonstrate an autocorrelation analysis of the spatially-resolved thickness measurements.

### Optimization at the stage of data acquisition

#### Influence of increased electron dose

We first investigated the influence of increasing electron dose, which has a favorable impact on contrast-to-noise ratio (CNR) but also produces damage in cryo-preserved samples (34). We acquired dose-fractionated movies of the same field of vesicles exposed continuously to the electron beam for 60 sec at a defocus of 2 μm. The impact of increasing accumulated dose on vesicle characteristics was then assessed by adjusting the number of frames included in the drift correction step to create the final images. This mode of analysis ensures that dose was the sole factor governing the subsequent comparison, as samples collected at different locations would be subject to other potentially complicating factors such as different ice thickness or astigmatism due to beam shifts. To eliminate contrast gradients across the images, data was high pass filtered before analysis. The impact of high pass filtering on measured membrane thickness and CNR were assessed and no differences were detected, as expected with a simple high pass filter. More direct comparisons on the impact of high pass filtering can be found in the section below comparing the influence of defocus on measured parameters (see Fig. 2).

As seen in Fig. 1A, increasing the electron dose has the expected impact on image CNR (here, for DOPC vesicles), which increases linearly up to ~ 40 e^−^/Å^2^ before leveling off. The gain in contrast is accompanied by a systematic decrease in the calculated trough-to-trough distance (D_TT_) of the vesicle bilayers as the sample accumulates damage. The impact of dose is even more dramatic in vesicles prepared from lipids that undergo liquid-liquid phase separation (Fig. 1B). For this analysis, the number of frames in each drift corrected image was normalized to avoid bias due to increased CNR. At the low dose of 13 e^−^/Å^2^ (Fig. 1B, green symbols) two distinct peaks are clearly observed in the D_TT_ histogram. These correspond to a thinner Ld phase with a mean thickness of ~ 30 Å coexisting with a thicker Lo phase of ~ 38 Å thickness, consistent with our previous report (30). Raising the dose to 39 e^−^/Å^2^ (Fig. 1B, magenta symbols) negatively impacts the ability to discriminate the Ld and Lo peaks. A further leftward shift of the peak and smearing of the distribution occurs at 52 and 65 e^−^/Å^2^ (Fig. 1B, blue and red symbols, respectively). The impacts of increasing dose on bilayer thickness was quantified and is shown in Table 1. The average D_TT_ of the ternary sample decreases with increasing dose, consistent with bilayer thinning resulting from beam-induced damage as seen in the DOPC sample. Representative images of the dose at 13, 39, and 65 e^−^/Å^2^ show detectable but subtle differences (Fig. 1, panels D-E, respectively).

**Figure 1.**
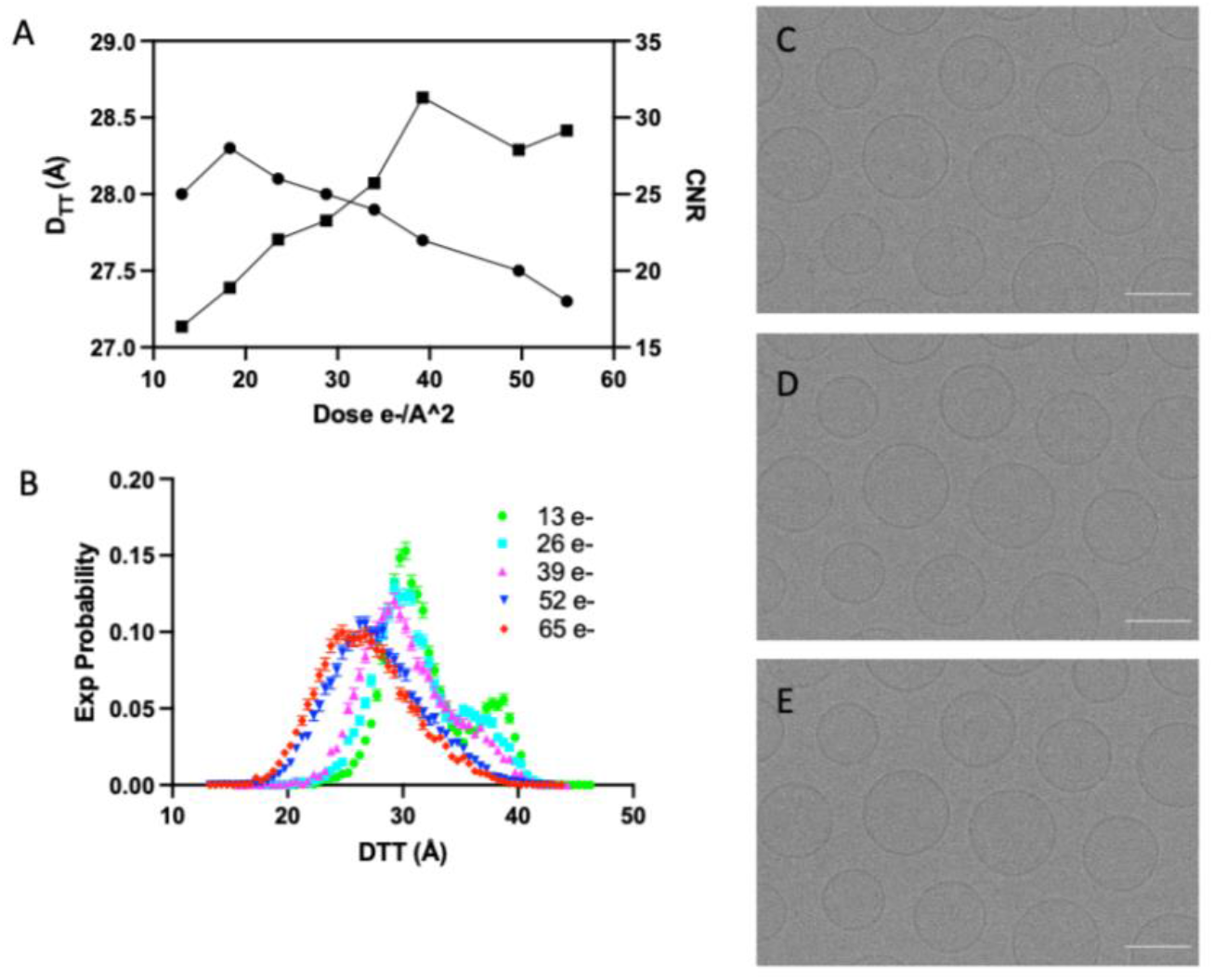
Influence of electron dose on bilayer thickness and image contrast. (A) Increasing the electron dose from 13-40 *e*^−^/Å^2^ results in an approximate doubling of the image contrast-to-noise ratio (CNR, squares) that is accompanied by a 2% decrease in the measured trough-trough distance D_TT_ (circles) in DOPC bilayers. A further increase in dose to 55 *e*^−^/Å^2^ causes an additional 2% thinning but without improvement in CNR. (B) In a phase-separated ternary mixture, two distinct peaks are observed in the D_TT_ histogram at low electron dose (green dots) corresponding to a thin Ld phase and thicker Lo phase. Upon increasing dose, the bimodal character of the distribution is lost and there is a leftward shift in the peak D_TT_ distribution. Data represents the average D_TT_ from ~125 vesicles from 5 separate images processed for dose as described in Methods. (C, D, E) Individual images from the data shown in Panel (B) from 13, 39 and 64 *e*^−^/Å^2^, respectively. Scale bars are 100 nm.

**Table 1.**
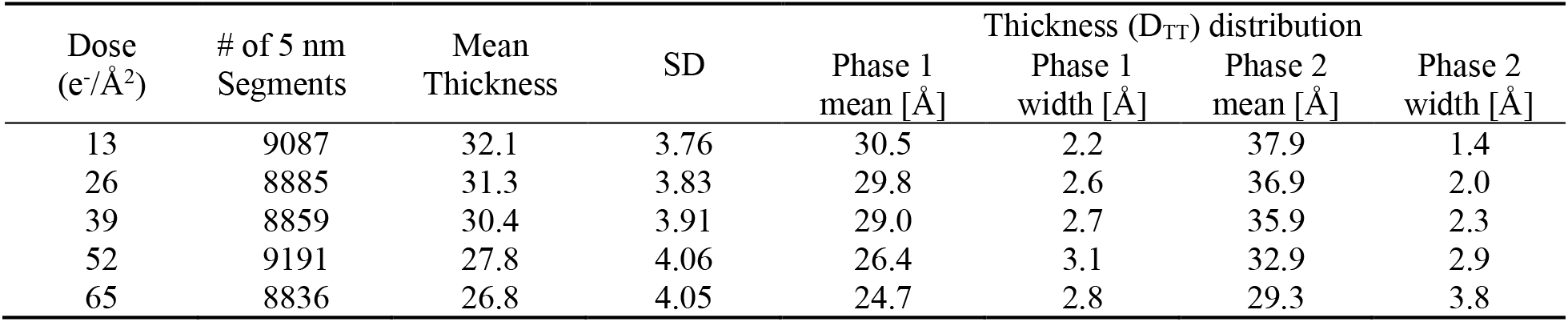
Impact of increasing dose on bilayer thickness.

| Dose<br>($e^-/\text{\AA}^2$) | # of 5 nm<br>Segments | Mean<br>Thickness | SD | Thickness ($D_{TT}$) distribution | | | |
| --- | --- | --- | --- | --- | --- | --- | --- |
| | | | | Phase 1<br>mean [$\text{\AA}$] | Phase 1<br>width [$\text{\AA}$] | Phase 2<br>mean [$\text{\AA}$] | Phase 2<br>width [$\text{\AA}$] |
| 13 | 9087 | 32.1 | 3.76 | 30.5 | 2.2 | 37.9 | 1.4 |
| 26 | 8885 | 31.3 | 3.83 | 29.8 | 2.6 | 36.9 | 2.0 |
| 39 | 8859 | 30.4 | 3.91 | 29.0 | 2.7 | 35.9 | 2.3 |
| 52 | 9191 | 27.8 | 4.06 | 26.4 | 3.1 | 32.9 | 2.9 |
| 65 | 8836 | 26.8 | 4.05 | 24.7 | 2.8 | 29.3 | 3.8 |

We conclude from this data that increasing dose has a favorable impact on CNR, but also results in specimen damage that negatively impacts the ability to measure bilayer thickness and thus to discriminate between phases of different inherent thickness. As evident in Fig. 1B, optimizing the dose becomes particularly important if the goal is to interrogate phase coexistence in individual vesicles. Having identified a total dose of ~ 20 e^−^/Å^2^ as a reasonable compromise between enhanced CNR and minimal impact on bilayer structure, all additional data was collected targeting this dose.

#### Influence of defocus

Because intensity in low-dose cryo-EM is generated primarily by phase contrast, defocusing has a large influence on image contrast. An example is shown in the series of micrographs of DOPC vesicles in Fig. 2A–C, in which defocus was varied from 0.95-3.3 μm: the enhancement of image contrast with increasing defocus is striking. To within measurement uncertainty, D_TT_ values do not depend on defocus in this set of images, as shown in the plot of DTT vs defocus (Fig. 2D). However, the variance in the measurements are smaller when defocus is centered around 2 μm. We also quantified the CNR as a function of defocus (Fig. 2E), revealing a steep positive correlation up to ~ 2-2.5 μm that is reversed with a further increase in defocus. We conclude that a defocus value close to 2 μm is optimal for contrast and membrane thickness quantification.

**Figure 2.**
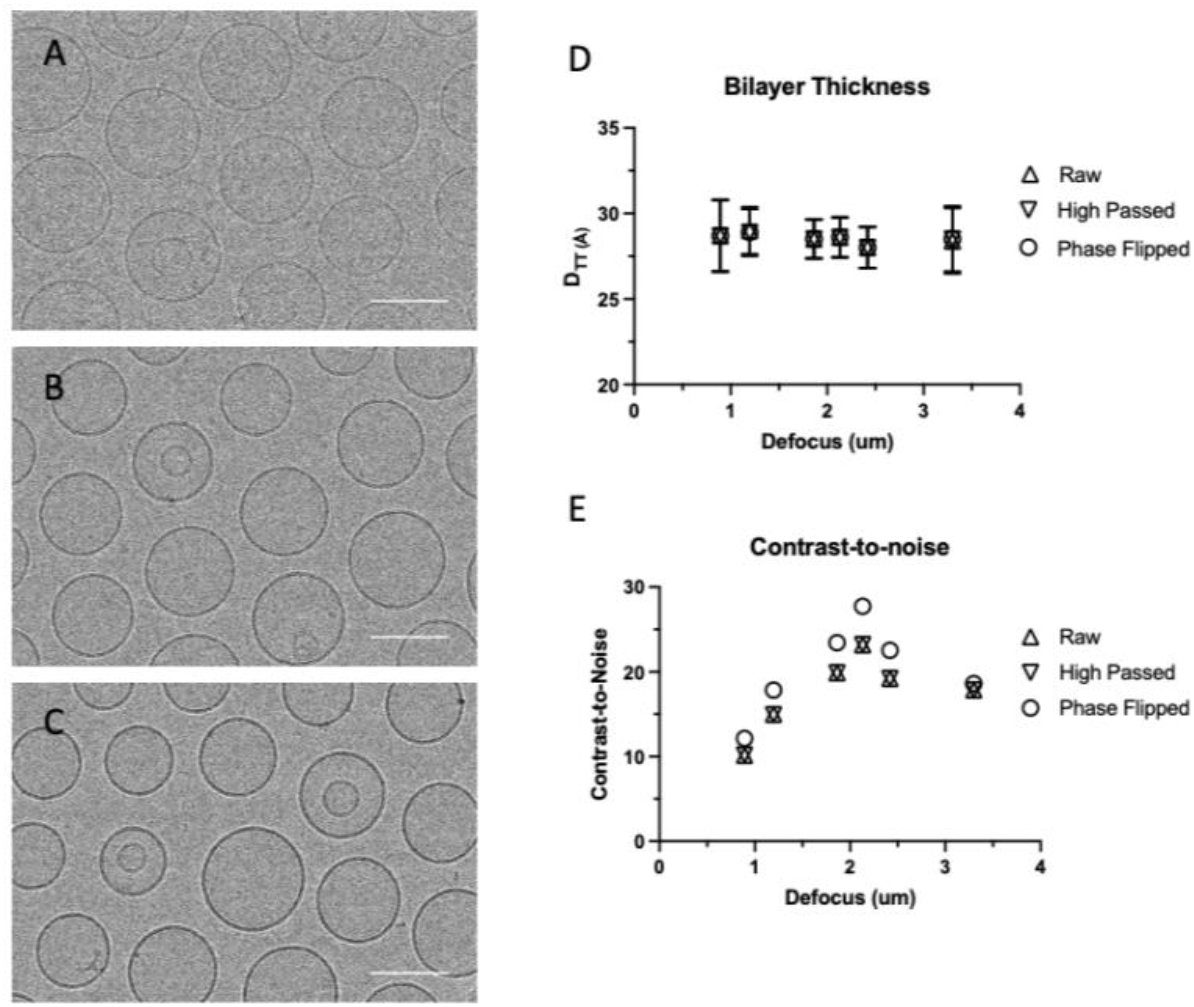
Influence of defocus on bilayer thickness and image contrast. (A, B, C) Representative sections from images of DOPC vesicles obtained at values of defocus from top to bottom: 0.95, 2.1, and 3.3 μm. Scale bars are 100 nm. (D) Average bilayer thickness D_TT_ quantified from 50-60 vesicles from images at the indicated defocus values, all collected at ~20 e^−^/Å^2^. For these sets of images, comparisons were made between raw data, high pass filtered data, and high-passed filtered and phase flipped data. There was no discernable difference in D_TT_ measured across defoci or among the filters applied. (E) Contrast-to-noise ratio (CNR) quantified from the same 50-60 vesicles each from the images analyzed in (D). There is an increase and then decrease in CNR at increasing values of defocus. Additionally, phase flipping of the data showed a modest but reproducible increase in CNR up to 2.2 μm defocus.

#### Influence of energy filtering

Energy filtering is a common strategy to improve signal-to-noise in cryo-EM by minimizing the contribution of inelastically scattered electrons in the final image. To evaluate the impact of energy filtering, we collected data from areas of the same EM grid of a DOPC sample, with and without energy filtering (20 eV slit in zero loss mode). Figure 3 shows representative images collected without (panel A) and with (panel B) energy filtering. The impact of energy filtering is most easily seen by comparing radially averaged vesicle intensity profiles of images collected at a similar defocus and dose, either without or with energy filtering as shown in Fig. 3C. The energy filtered profiles (Fig. 3C, green profiles) show less intensity variation than unfiltered profiles (Fig. 3C, magenta profiles). Despite the reduction in intensity variation, when data from 80-120 vesicles across five separate images (matched for defocus between 1.8-2.0 μm) are pooled, there is little discernible difference in the peak position or width of the thickness probability distribution of energy filtered vs. unfiltered data (Fig. 3D and Table 2). Additionally, there is no difference in CNR (Table 2). We conclude that collecting data using an energy filter offers a discernable but minor advantage for measurements of bilayer thickness distributions. Further, we repeated this analysis of CNR and bilayer thickness in phase-separated vesicles with and without energy filtering (Fig. 4). Energy filtering results in a small shift in the apparent phase fractions toward the thinner disordered phase and a slight rightward shift in the thickness distribution (Fig. 4C, Table 2). We conclude that energy filtering offers no distinct advantages for the analyses used here. Instead, careful attention to dose and defocus are more important considerations for optimizing images for membrane thickness measurements.

**Figure 3.**
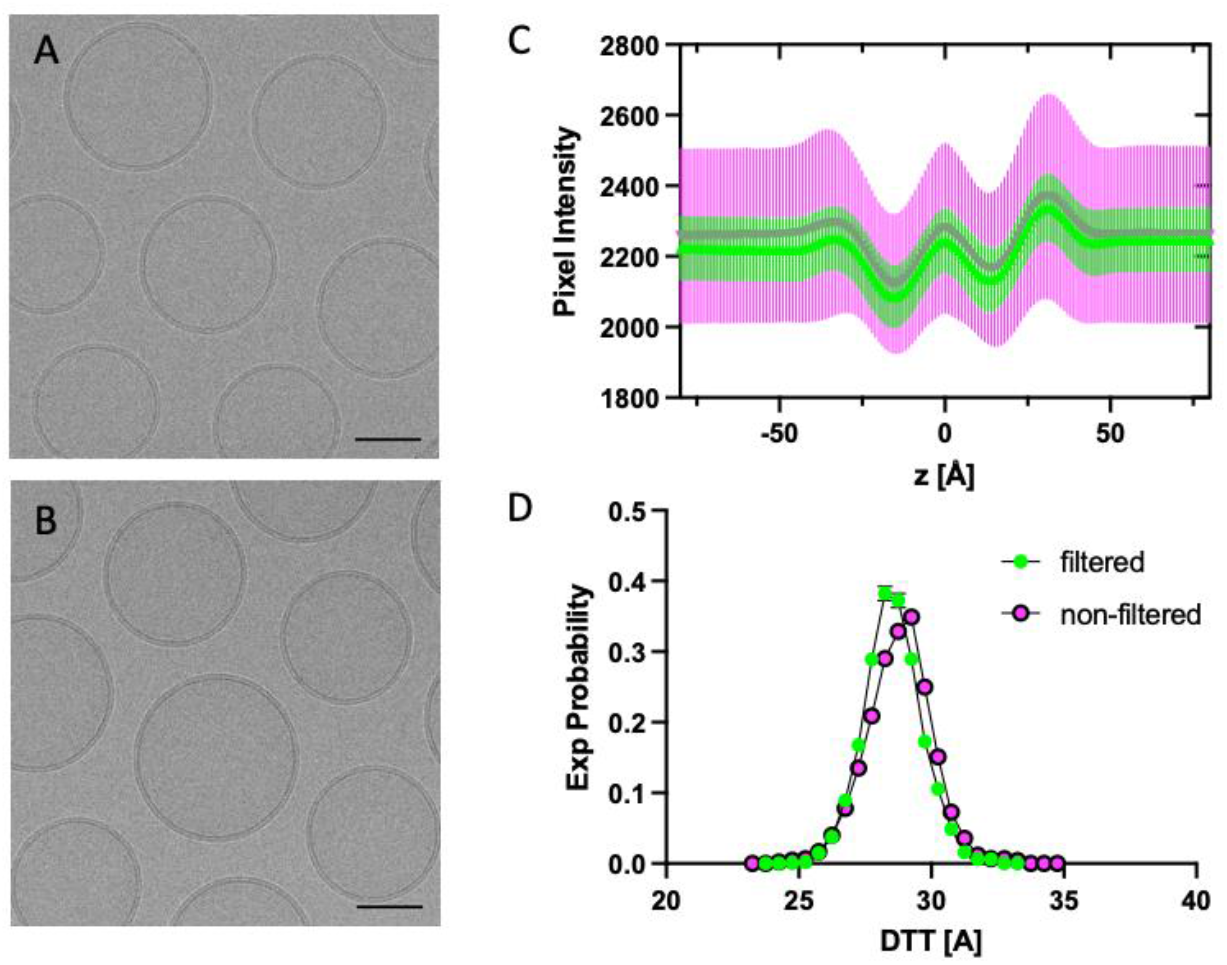
Influence of energy filtering on bilayer thickness and image contrast. Representative images of DOPC vesicles without (panel A) and with (panel B) energy filtering. Data collected at ~20 e^−^/Å^2^ and 2 um under focus. Scale bar is 50 nm. (C) Radially averaged intensity profiles for DOPC vesicles without (magenta) and with (green) energy filtering. (D) Bilayer thickness (D_TT_) histograms of images acquired with energy-filtered (green) or without energy filtered (magenta).

**Figure 4.**
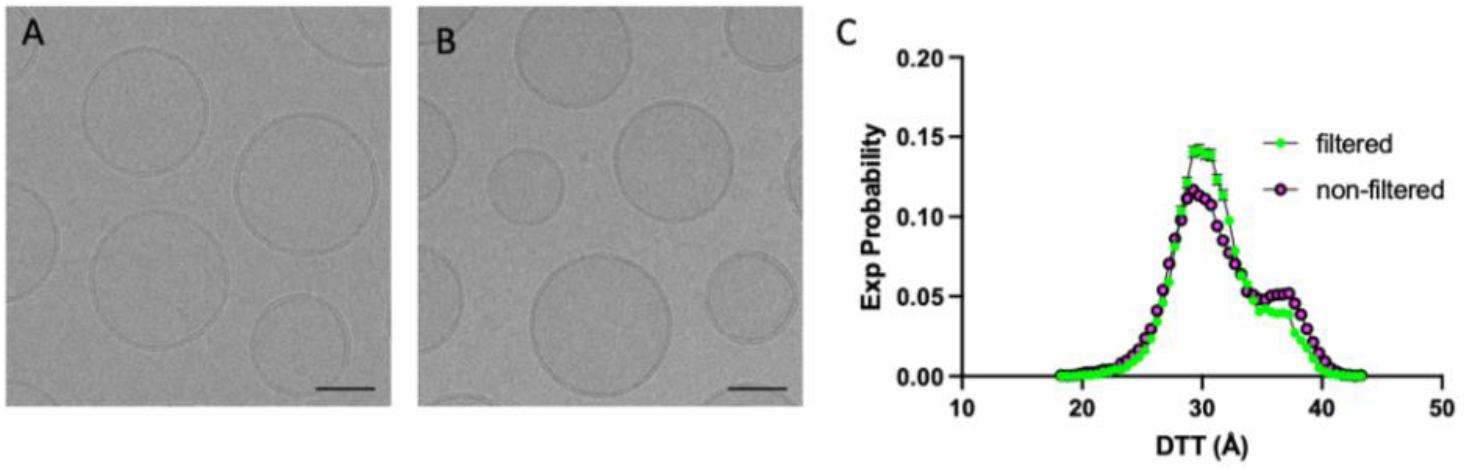
Influence of energy filtering on bilayer thickness and image contrast. Representative images of DPPC/DOPC/cholesterol vesicles without (panel A) and with (panel B) energy filtering. Data collected at ~20 e^−^/Å^2^ and 2 um under focus. Scale bar is 50 nm. (C) Bilayer thickness (D_TT_) histograms of images acquired with energy-filtered (green) or without energy filtered (magenta).

**Table 2.** Influence of energy filtering on bilayer thickness and image contrast.

| Sample | Energy filtered | N | CNR | Thickness ($D_{\text{TT}}$) distribution | | | |
| --- | --- | --- | --- | --- | --- | --- | --- |
| | | | | Phase 1 mean [$\text{\AA}$] | Phase 1 width [$\text{\AA}$] | Phase 2 mean [$\text{\AA}$] | Phase 2 width [$\text{\AA}$] |
| DOPC | No | 120 | 19.9 | 28.8 | 1.2 | -- | -- |
| DOPC | Yes | 86 | 20.7 | 28.5 | 1.1 | -- | -- |
| Ternary | No | 75 | 16.9 | 29.3 | 1.8 | 35.3 | 2.0 |
| Ternary | Yes | 91 | 17.0 | 30.3 | 2.4 | 36.6 | 1.2 |

### 3.2 Optimization of image post-processing and analysis

#### Influence of phase-flipping and high pass filtering

Imaging strives to provide as accurate a reproduction of the original object as possible. Achieving this goal is particularly challenging in low dose cryo-EM because signal is largely (> 90%) from phase-contrast that is dependent on a number of instrument properties, including accelerating voltage and spherical aberrations (which are largely fixed) and the setting of defocus as highlighted above. A striking example of the impact of defocus on image formation is evident in Fig. 2, which clearly shows how increasing defocus increases image contrast. This effect, termed the contrast transfer function (CTF), has an oscillating behavior where the information transferred into the image is a complex function dependent on the frequency of spatial information (Fig. 5). Additional factors such as the envelope function of the microscope tend to selectively dampen the transmission of higher frequency information (Fig. 5). These well-described phenomena are common to all cryo-EM studies, and image restoration algorithms have been developed to recover (as best as possible) information in the digital image to accurately represent the original object. The simplest way to correct for CTF artifacts is to invert the sign of the phases in the negative regions (“phase-flipping”) after estimating the image’s CTF, an operation that is typically performed in conjunction with removing frequencies near the zeros of the CTF that contain essentially no structural information and contribute only to noise.

**Figure 5.**
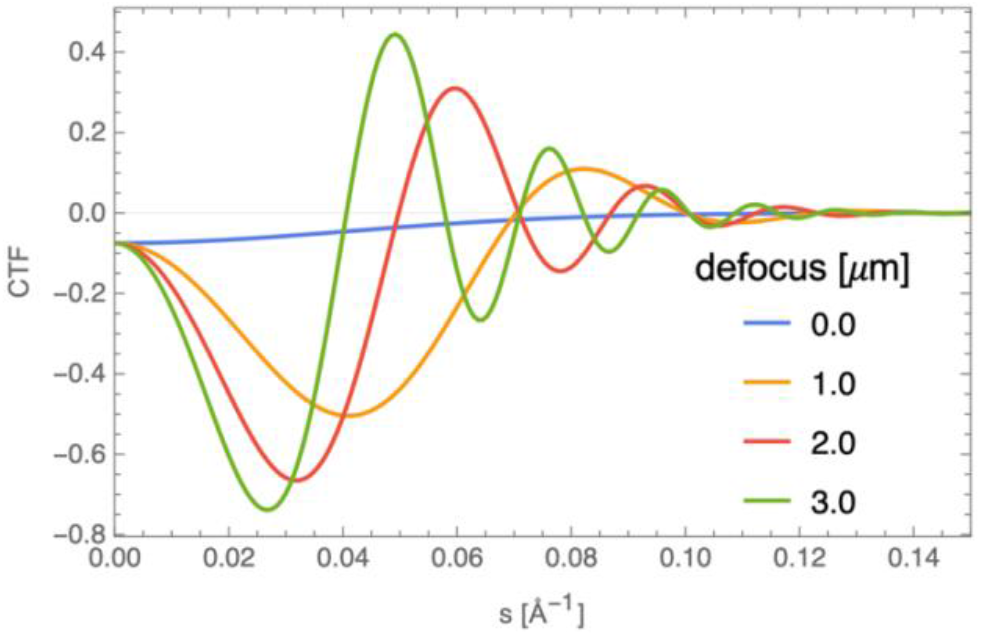
Influence of defocus on the contrast transfer function. The CTF given by Eqs. 1–2 (Methods) was simulated at various levels of defocus. The amplitude and frequency of the oscillations increases with increasing defocus: Δ*Z* = 0.0 (blue), 1.0 (orange), 2.0 (red) and 3.0 (green). The decreased amplitude of the CTF at higher spatial frequencies is the result of the envelope function of the microscope (the exponential term in Eq. 1). Values of the fixed parameters were: *Q* = 7.5%, *B* = 300 Å^2^, *λ* = 0.0197 Å, *C*_*s*_ = 0.

To assess the impact of these post-processing steps, we selected micrographs that fell within a narrow range of defocus (1.8-2.0 μm) for analysis. Images were typically high pass filtered to remove low frequency variation in intensity that can arise from sources like gradients in ice thickness across the field of view. Figure 6 shows representative images before (panel A) and after (panel B) processing by a high pass filter either alone or with phase-flipping (panel C). There is little visually identifiable difference in images before or after these processing steps. Nevertheless, quantitative image analysis revealed two impacts of phase-flipping. First, there was a small but consistent increase in the D_TT_ values of phase-flipped data, independent of energy filtering (Fig. 6D and 6E, Table 3). Second, there was a consistent increase in the CNR of phase-flipped images (Table 3). In contrast, high pass filtering had essentially no impact on CNR or measured bilayer thickness (Fig. 2B and C, Table 3). We conclude that phase-flipping has a favorable impact on the quantification of membrane thickness, presumably due to increases CNR, and should be considered when optimizing analysis of membranes using cryo-EM.

**Figure 6.**
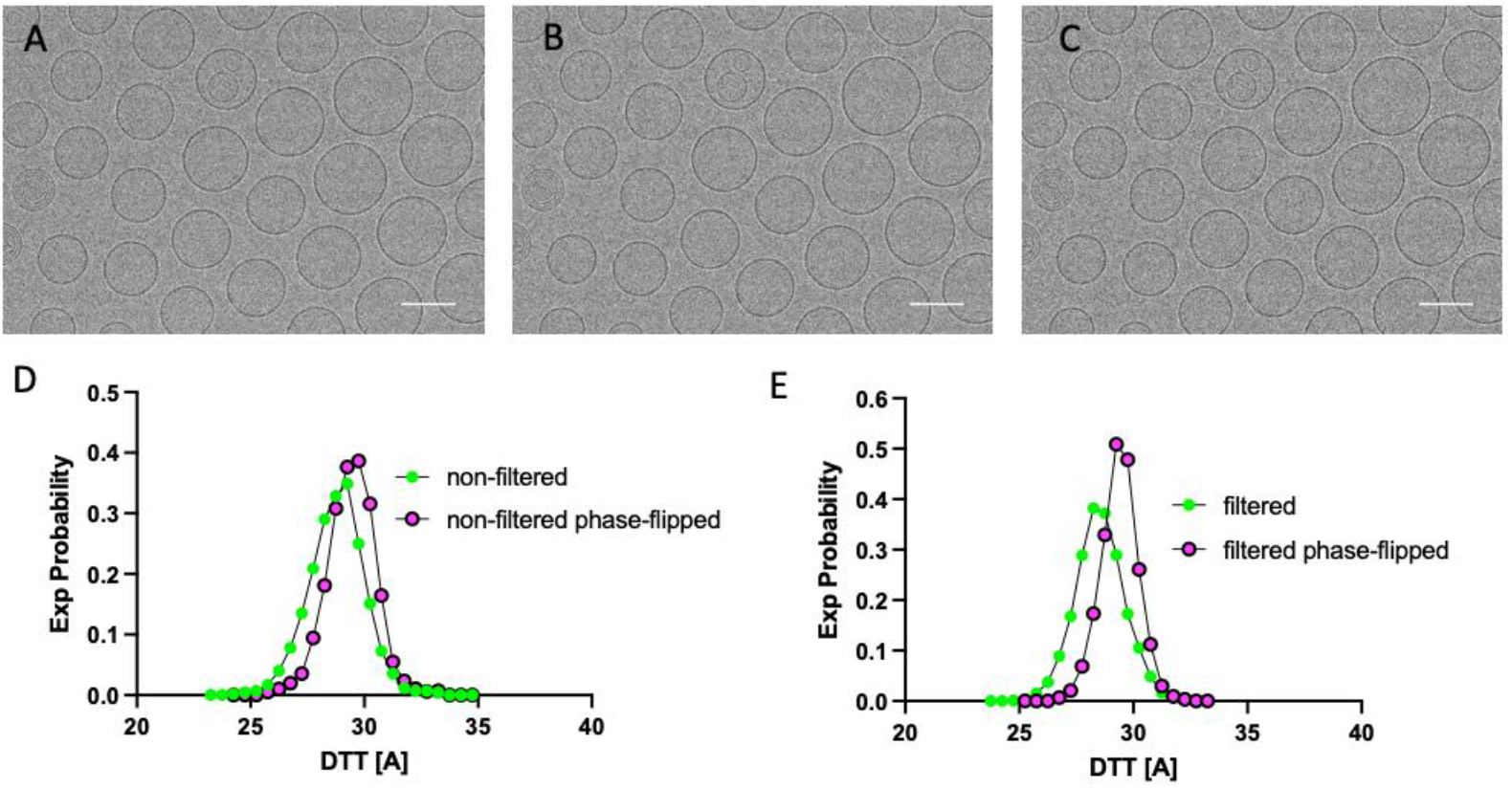
Influence of phase-flipping on bilayer thickness measurements. (a) Representative image of DOPC vesicles before (A) and after application of high pass filter separately (B) and with phase-flipping (C). Scale bars are 100 nm. Also shown are bilayer thickness (D_TT_) histograms of images before (green) and after (magenta) phase-flipping for non-energy filtered (D) and energy-filtered (E) images.

**Table 3.**
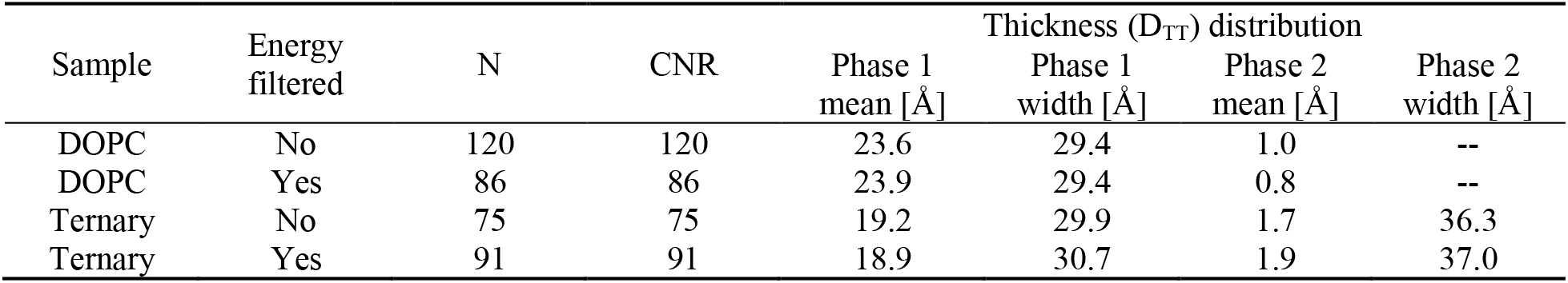
Influence of phase-flipping on bilayer thickness and image contrast.

| Sample | Energy filtered | N | CNR | Thickness ($D_{TT}$) distribution | | | |
| --- | --- | --- | --- | --- | --- | --- | --- |
| | | | | Phase 1 mean [$\text{\AA}$] | Phase 1 width [$\text{\AA}$] | Phase 2 mean [$\text{\AA}$] | Phase 2 width [$\text{\AA}$] |
| DOPC | No | 120 | 120 | 23.6 | 29.4 | 1.0 | -- |
| DOPC | Yes | 86 | 86 | 23.9 | 29.4 | 0.8 | -- |
| Ternary | No | 75 | 75 | 19.2 | 29.9 | 1.7 | 36.3 |
| Ternary | Yes | 91 | 91 | 18.9 | 30.7 | 1.9 | 37.0 |

#### Autocorrelation of spatially resolved thickness to assess lateral heterogeneity

We previously showed that coexistence of ordered and disordered phases can be inferred from histograms of spatially resolved bilayer thickness (D_TT_) measurements (Heberle et al., 2020). Histograms of D_TT_ measured at 5 nm resolution along the projected vesicle circumference show a single peak for DOPC vesicles that is well fit by a single Gaussian distribution (e.g., Fig. 3D, Fig. 5D and E). In contrast, histograms from ternary vesicles show a bimodal thickness distribution (Fig. 1B and 4C).

Because the ability to discriminate two distinct thickness populations depends on the intrinsic thickness difference between the coexisting phases and likely also on domain size, we investigated an alternative approach to identifying lateral heterogeneity through the spatial autocorrelation of D_TT_ calculated for individual vesicles and averaged over the population of vesicles. Figure 7 shows the resulting autocorrelation data for DOPC and ternary vesicles with (green) and without (magenta) energy filtering. For DOPC vesicles, the autocorrelation curves are essentially structureless as expected for a uniform bilayer. There is no discernible heterogeneity in the thickness of the membrane across length scales. In contrast, curves for ternary vesicles show a strong positive thickness correlation at short lag distances that gradually decays to a positive baseline value at a distance of 20-30 nm (Fig. 7, light blue triangles). A plausible explanation for the positive baseline value at longer lags is the presence of vesicles that exhibit only Ld or Lo in projection. The structure in the autocorrelation data from the ternary mixture is consistent with lateral heterogeneity and may provide an important tool for assessing the presence of phase separation in cases of smaller intrinsic thickness differences between coexisting phases. As noted above, for both types of vesicles, energy filtering has practically no effect on the autocorrelation curves.

**Figure 7.**
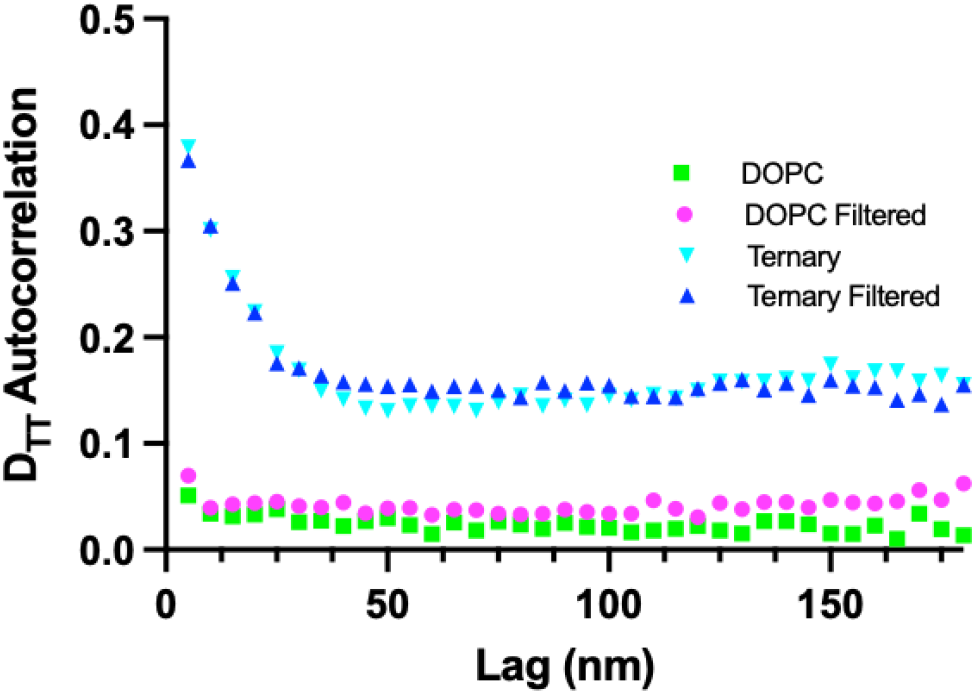
Spatial autocorrelation of bilayer thickness. Spatial autocorrelation of D_TT_ for DOPC (green and magenta) and ternary vesicles (light blue and dark blue) with (magenta and dark blue) and without (green and light blue) energy filtering.

## Conclusion

Our results provide concrete guidelines for minimizing artifacts while maximizing the information content of cryo-EM images of lipid bilayers. Specifically, total dose should be kept below 20 e^−^/Å^2^ to minimize specimen damage that negatively impacts the ability to measure membrane thickness. Additionally, ~ 2 μm underfocus was found to provide good contrast while avoiding smearing that leads to artifacts in assessing membrane thickness. Energy filtering of the primary data did not provide any distinct advantages for assessing membrane characteristics. Image processing using high pass filtering and phase-flipping were found to increase the quality and contrast-to-noise ratio in the data and are recommended for optimizing analyses of lipid bilayer thickness. Finally, we demonstrate the use of a spatial autocorrelation as an additional tool to assess the presence of phase separation. Autocorrelation curves may prove useful for mixtures with a smaller intrinsic thickness difference between coexisting phases, or with inherently smaller domains that may be difficult to see by eye.

## Author contributions

F.A.H. and M.N.W. designed the research. D.W., H.L.S., and M.N.W. conducted experiments. D.W., H.L.S., F.A.H., and M.N.W. analyzed data. F.A.H. and M.N.W. wrote the article. D.W., H.L.S., F.A.H., and M.N.W. revised the manuscript.

## Acknowledgements

We acknowledge the help of Mr. Venkata Mallampalli and Dr. Guizhen Fan for data collection on the Krios microscope supported through the McGovern Medical School Structural Biology Center. We also are grateful to Dr. Tristan Bepler for useful discussions and for providing the machine learning algorithm, MEMNET, for automated contouring of vesicles. Finally, we are grateful to Mr. Dustin Morado for numerous helpful discussions on image processing and, in particular helping with routines for CTF estimation and phase-flipping. This work was supported by NSF Grant MCB-1817929 (to F.A.H.), NSF Grant CHE-220412 (to F.A.H and M.N.W.) and NIH Grant R01GM138887 (to F.A.H and M.N.W.). M.N.W. acknowledges the William Wheless III Professorship.

## References

1. Lorent, J.H., K.R. Levental, L. Ganesan, G. Rivera-Longsworth, E. Sezgin, M. Doktorova, E. Lyman, and I. Levental. 2020. Plasma membranes are asymmetric in lipid unsaturation, packing and protein shape. Nature Chemical Biology. 16:644–652.

2. Sezgin, E., I. Levental, S. Mayor, and C. Eggeling. 2017. The mystery of membrane organization: Composition, regulation and roles of lipid rafts. Nature Reviews Molecular Cell Biology. 18:361–374.

3. Simons, K., and E. Ikonen. 1997. Functional rafts in cell membranes. Nature. 387:569–572.

4. Lingwood, D., and K. Simons. 2010. Lipid rafts as a membrane-organizing principle. Science (New York, N.Y.). 327:46–50.

5. Pierce, S.K. 2002. Lipid rafts and B-cell activation. Nature reviews. Immunology. 2:96–105.

6. Heberle, F.A., and G.W. Feigenson. 2011. Phase separation in lipid membranes. Cold Spring Harbor Perspectives in Biology. 3:1–13.

7. Hjort Ipsen, J., G. Karlström, O.G. Mourtisen, H. Wennerström, and M.J. Zuckermann. 1987. Phase equilibria in the phosphatidylcholine-cholesterol system. Biochimica et biophysica acta. 905:162–172.

8. Veatch, S.L., P. Cicuta, P. Sengupta, A. Honerkamp-Smith, D. Holowka, and B. Baird. 2008. Critical fluctuations in plasma membrane vesicles. ACS chemical biology. 3:287–293.

9. Chiang, Y.W., J. Zhao, J. Wu, Y. Shimoyama, J.H. Freed, and G.W. Feigenson. 2005. New method for determining tie-lines in coexisting membrane phases using spin-label ESR. Biochimica et biophysica acta. 1668:99–105.

10. Feigenson, G.W., and J.T. Buboltz. 2001. Ternary phase diagram of dipalmitoyl-PC/dilauroyl-PC/cholesterol: nanoscopic domain formation driven by cholesterol. Biophysical journal. 80:2775–2788.

11. Heberle, F.A., J.T. Buboltz, D. Stringer, and G.W. Feigenson. 2005. Fluorescence methods to detect phase boundaries in lipid bilayer mixtures. Biochimica et Biophysica Acta – Molecular Cell Research. 1746:186–192.

12. Pathak, P., and E. London. 2011. Measurement of lipid nanodomain (raft) formation and size in sphingomyelin/POPC/cholesterol vesicles shows TX-100 and transmembrane helices increase domain size by coalescing preexisting nanodomains but do not induce domain formation. Biophysical journal. 101:2417–2425.

13. Heberle, F.A., M. Doktorova, S.L. Goh, R.F. Standaert, J. Katsaras, and G.W. Feigenson. 2013. Hybrid and nonhybrid lipids exert common effects on membrane raft size and morphology. Journal of the American Chemical Society. 135:14932–14935.

14. Marquardt, D., F.A. Heberle, J.D. Nickels, G. Pabst, and J. Katsaras. 2015. On scattered waves and lipid domains: Detecting membrane rafts with X-rays and neutrons. Soft Matter. 11:9055–9072.

15. Khadka, N.K., C.S. Ho, and J. Pan. 2015. Macroscopic and Nanoscopic Heterogeneous Structures in a Three-Component Lipid Bilayer Mixtures Determined by Atomic Force Microscopy. Langmuir : the ACS journal of surfaces and colloids. 31:12417–12425.

16. Moss, F.R., and S.G. Boxer. 2016. Atomic Recombination in Dynamic Secondary Ion Mass Spectrometry Probes Distance in Lipid Assemblies: A Nanometer Chemical Ruler. Journal of the American Chemical Society. 138:16737–16744.

17. Wilson, R.L., J.F. Frisz, H.A. Klitzing, J. Zimmerberg, P.K. Weber, and M.L. Kraft. 2015. Hemagglutinin clusters in the plasma membrane are not enriched with cholesterol and sphingolipids. Biophysical journal. 108:1652–1659.

18. Sezgin, E. 2017. Super-resolution optical microscopy for studying membrane structure and dynamics. Journal of physics. Condensed matter : an Institute of Physics journal. 29.

19. Goksu, E.I., and M.L. Longo. 2010. Ternary lipid bilayers containing cholesterol in a high curvature silica xerogel environment. Langmuir : the ACS journal of surfaces and colloids. 26:8614–8624.

20. Gunderson, R.S., and A.R. Honerkamp-Smith. 2018. Liquid-liquid phase transition temperatures increase when lipid bilayers are supported on glass. Biochimica et biophysica acta. Biomembranes. 1860:1965–1971.

21. Owen, D.M., A. Magenau, D.J. Williamson, and K. Gaus. 2013. Super-resolution imaging by localization microscopy. Methods in molecular biology (Clifton, N.J.). 950:81–93.

22. Chorev, D.S., and C.V. Robinson. 2020. The importance of the membrane for biophysical measurements. Nature chemical biology. 16:1285–1292.

23. Egelman, E.H. 2016. The Current Revolution in Cryo-EM. Biophysical journal. 110:1008–1012.

24. Kühlbrandt, W. 2014. Cryo-EM enters a new era. eLife. 3:e03678.

25. Gipson, P., Y. Fukuda, R. Danev, Y. Lai, D.H. Chen, W. Baumeister, and A.T. Brunger. 2017. Morphologies of synaptic protein membrane fusion interfaces. Proceedings of the National Academy of Sciences of the United States of America. 114:9110–9115.

26. Han, M., Y. Mei, H. Khant, and S.J. Ludtke. 2009. Characterization of antibiotic peptide pores using cryo-EM and comparison to neutron scattering. Biophysical journal. 97:164–172.

27. Tahara, Y., and Y. Fujiyoshi. 1994. A new method to measure bilayer thickness: cryo-electron microscopy of frozen hydrated liposomes and image simulation. Micron (Oxford, England : 1993). 25:141–149.

28. Wang, L., P.S. Bose, and F.J. Sigworth. 2006. Using cryo-EM to measure the dipole potential of a lipid membrane. Proceedings of the National Academy of Sciences of the United States of America. 103:18528–18533.

29. Cornell, C.E., A. Mileant, N. Thakkar, K.K. Lee, and S.L. Keller. 2020. Direct imaging of liquid domains in membranes by cryo-electron tomography. Proceedings of the National Academy of Sciences of the United States of America. 117:19713–19719.

30. Heberle, F.A., M. Doktorova, H.L. Scott, A.D. Skinkle, M.N. Waxham, and I. Levental. 2020. Direct label-free imaging of nanodomains in biomimetic and biological membranes by cryogenic electron microscopy. Proceedings of the National Academy of Sciences of the United States of America. 117:19943–19952.

31. Mastronarde, D.N. 2005. Automated electron microscope tomography using robust prediction of specimen movements. Journal of structural biology. 152:36–51.

32. Zheng, S.Q., E. Palovcak, J.P. Armache, K.A. Verba, Y. Cheng, and D.A. Agard. 2017. MotionCor2: anisotropic correction of beam-induced motion for improved cryo-electron microscopy. Nature methods. 14:331–332.

33. Scott, H.L., A. Skinkle, E.G. Kelley, M.N. Waxham, I. Levental, and F.A. Heberle. 2019. On the Mechanism of Bilayer Separation by Extrusion, or Why Your LUVs Are Not Really Unilamellar. Biophysical Journal. 117.

34. Baker, L.A., and J.L. Rubinstein. 2010. Radiation damage in electron cryomicroscopy. Methods in enzymology. 481:371–388.

